# Detecting and Subtyping Ketoacidosis from Metabolomic Patterns in Forensic Casework

**DOI:** 10.64898/2026.03.09.710563

**Authors:** Ralph E. C. Monte, Rasmus Magnusson, Carl Söderberg, Henrik Gréen, Albert Elmsjö, Elin Nyman

## Abstract

Subtyping of ketoacidosis, a metabolic state characterized by blood acidification due to various causes, remains challenging in forensic casework. Postmortem omics samples paired with machine learning offers an independent tool to address this challenge. However, such data, especially related to real forensic cases, are rare. In Sweden, high-resolution mass spectrometry data routinely collected in forensic toxicology, can be leveraged for metabolomic analysis. Here, we integrate postmortem metabolomics and machine learning models to detect and subtype ketoacidosis-related deaths using real forensic cases in Sweden. From femoral blood samples of 109 alcoholic ketoacidosis cases, 220 diabetic ketoacidosis cases, 140 hypothermia cases, and 1,229 controls (hanging cases), we developed and tested three machine learning models, which achieved over 90% accuracy in ketoacidosis detection and over 80% in subtyping. Validation with independent cohorts (21 starvation cases, 29 alcoholic controls, and 40 diabetic controls) confirmed robustness with over 80% of starvation cases classified as ketoacidosis-related. Feature clustering highlighted metabolites such as cortisol to be important for subtyping. In summary, our findings demonstrate that combining machine learning with postmortem metabolomics enables accurate detection and subtyping of ketoacidosis-related deaths, which is useful for forensic casework.

## Introduction

Cause of death determination in forensic casework can be challenging, particularly when different conditions have shared pathophysiological mechanisms. This complexity is evident in deaths caused by ketoacidosis. Ketoacidosis occurs during periods of decreased carbohydrate levels or availability, where ketone bodies accumulate due to increased lipolysis of free fatty acids [1]. This accumulation of ketone bodies leads to high anion gap metabolic acidosis and eventual death if untreated [2]. Ketoacidosis can occur due to several reasons, including diabetes (diabetic ketoacidosis, DKA), alcoholism (alcoholic ketoacidosis, AKA), hypothermia, and starvation [3–5]. In suspected ketoacidosis-related deaths, additional pathological findings are needed to determine cause of death, including forensic screening of the femoral blood to measure one or several ketone bodies and other biomarkers, like glucose [6,7]. In current forensic pathology practice, biomarkers are measured in vitreous humour due to its high stability postmortem [8–10]. However, these targeted measurements might not capture the broader metabolic context needed to differentiate between deaths with closely-related pathophysiologies, such as in the case of different types of ketoacidosis. This underlines the need of an objective, high-throughput approach that can provide a more comprehensive view of the metabolic state at the time of death.

As ketoacidosis is a metabolic condition, metabolomics data is a primary candidate for such an approach. Metabolomics is the qualitative and/or quantitative analysis of low molecular weight metabolites, which reflects the metabolic state of an organism at a certain time point [11,12]. These metabolite profiles can be compared across conditions to quantify the effects of treatments, physiological perturbations, or disease states [13]. In forensic science, postmortem metabolomics have emerged as a tool to, e.g., determine cause or time of death [14–16], or discovery of biomarkers related to drug abuse [17,18] or poisonings [19]. Yet, using metabolomics data for such analyses is non-trivial, partly due to its high-dimensional and complex nature. Thus, proper statistical tools are needed [20].

Machine learning (ML) can make use of such high-dimensional and complex data. ML methods can learn and identify patterns in the data that cannot be found with other, non-ML methods. ML methods can learn and identify patterns in the data [22] that cannot be found with other, non-ML methods [23]. ML integration with metabolomics has exponentially increased in the last 30 years, which highlights the interest and importance of integrating ML with metabolomics workflow [24]. Increasingly, postmortem metabolomics and a wide range of ML methods have been used to answer forensic questions [25]. For example, the postmortem interval (PMI) has been accurately predicted, employing a wide range of ML methods, like neural networks, random forest (RF) models, penalized linear regression [15,23,26]. Similarly, ML methods have been used as classification models for cause of death determination [27,28], introducing a more objective component into the forensic workflow. To the best of our knowledge, no study so far addresses the integration of ML methods and postmortem metabolomics to detect and subtype ketoacidosis-related deaths in real casework scenarios.

Nevertheless, Swedish forensic routines regularly collect postmortem metabolomics data, with a yearly output of approximately 5,500 real forensic cases. As such, femoral blood samples from deceased people routinely undergo toxicological screening using mass spectrometry ultra high-performance liquid chromatography quadrupole time of flight (UHPLC-QTOF) [21]. With the same method, endogenous metabolites are captured. This data, together with the forensic case-reports, presents an excellent opportunity for using ML to detect ketoacidosis-related deaths, and disentangle subtypes.

Here, we compiled a unique Swedish cohort of real forensic cases in which metabolomics data had been routinely collected from femoral blood during toxicological screening. From this resource, we train and evaluate supervised machine⍰learning models for classifying ketoacidosis and related conditions. Our approach included both binary and multinomial classification frameworks to assess the potential for detecting ketoacidosis and distinguishing its subtypes. Overall, this study aims to demonstrate how integrating supervised machine learning with postmortem metabolomics can improve the identification and characterization of ketoacidosis⍰related deaths, highlighting the broader potential of such methods in forensic practice.

## Results

We compiled and analysed postmortem metabolomics data from 1,788 femoral blood samples collected during forensic investigations between 2017 and 2020. These samples represented four groups used for model development: alcoholic ketoacidosis (AKA, n = 109), diabetic ketoacidosis (DKA, n = 220), hypothermia (n = 140), and controls (hanging cases) (*n* = 1,229) (Figure 1B). AKA, DKA, and hypothermia grouped together represent ketoacidosis cases (n = 469). Hanging cases were used as control cases since the cause of death for hanging cases is in general certain and hanging is assumed to cause minimal physiological alterations due to the rapid process of death. We also included an independent cohort used for model validation with three groups: starvation (n = 21), alcoholic controls (AC) (n = 29), and diabetic controls (DC) (*n* = 40) (Figure 2B). The characteristics of all groups are summarised in Table 1. In total, 4,484 metabolic features were identified after preprocessing the UHPLC-QTOF data from the included cases in this study.

**Figure 1.**
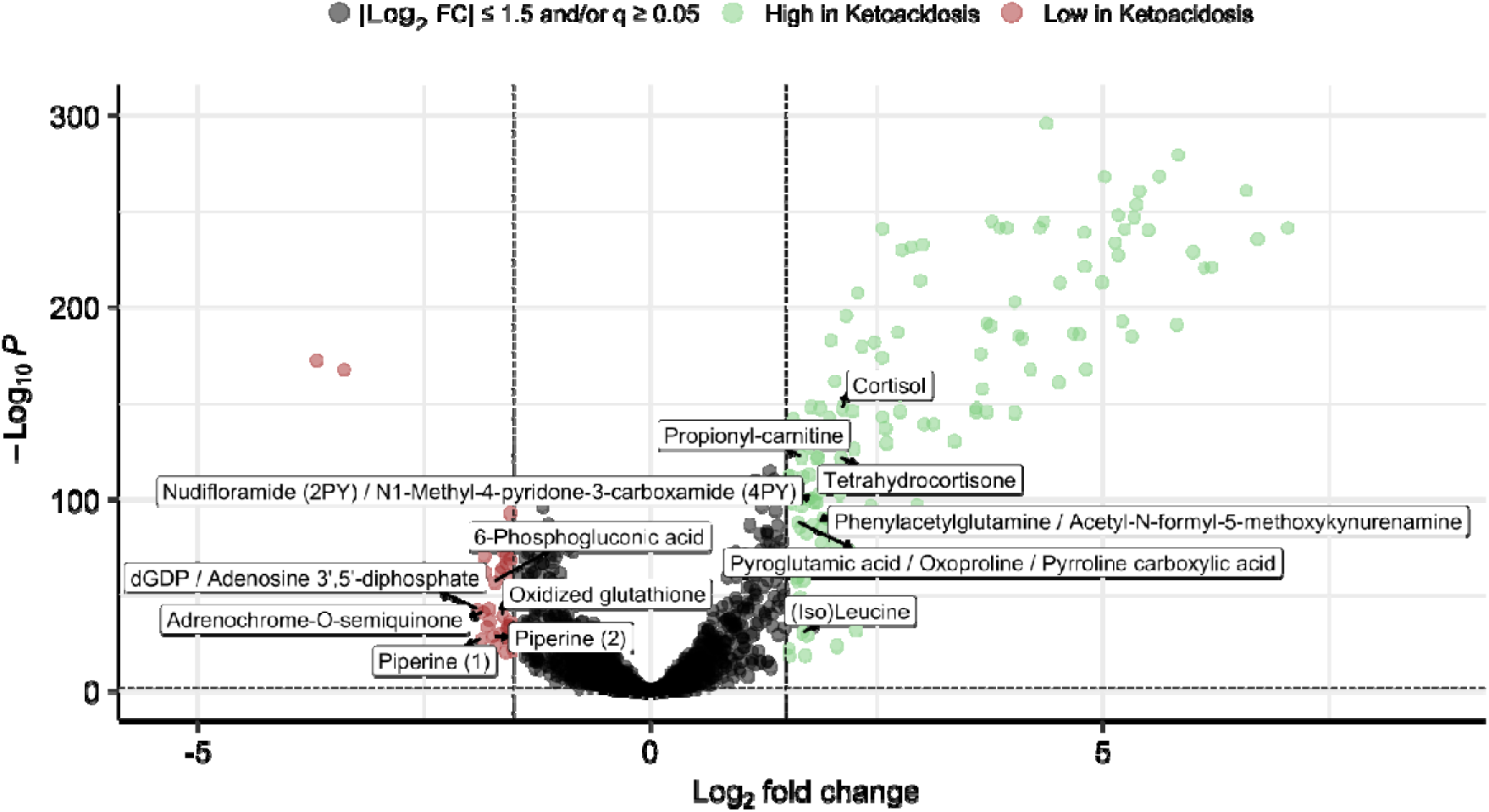
Volcano plot of the differences in abundances of metabolic features between ketoacidosis cases and controls. Metabolic features with a q-value of less than 0.05 and an absolute log_2_ fold change (Log_2_ FC) of greater than 1.5, i.e., significantly abundant metabolic features, were coloured red (lower in ketoacidosis cases: 37 metabolic features) or green (higher in ketoacidosis cases: 133 metabolic features). Significant metabolites that could be annotated are labelled.

**Figure 2.**
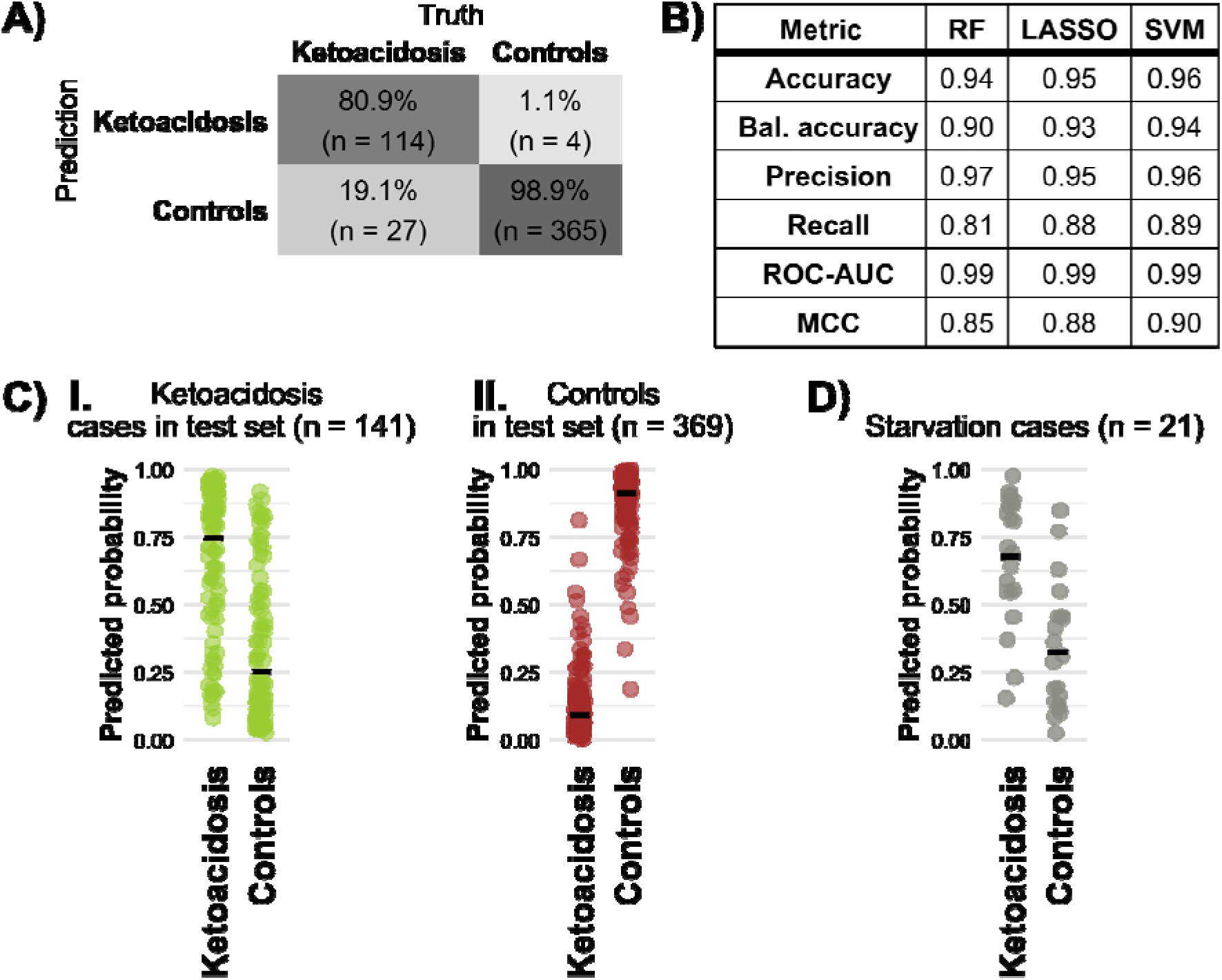
Supervised ML models could distinguish between ketoacidosis and controls. A) Confusion matrix for the binary RF model. B) Binary classification performance metrics of the three different binary models. Bal. accuracy = balanced accuracy. ROC-AUC = Receiver Operating Characteristic Area Under the Curve. MCC = Matthews correlation coefficient. C) Predicted probabilities of different classifications of test set data by the binary RF model. I: ketoacidosis cases, *n* = 141. II: controls, *n* = 369. Means denoted with black bars. D) Predicted probabilities of different classifications by the binary RF model of the starvation cases. Means denoted with black bars. *Note: a predicted probability > 0.5 leads to classification as ketoacidosis, while a predicted probability of* ≤ *0.5 leads to a classification as controls*.

**Table 1.** Demographic overview of the included groups.

| | AKA<br>($n = 109$) | DKA<br>($n = 220$) | Hypothermia<br>($n = 140$) | Hanging<br>($n = 1,229$) | Starvation<br>($n = 21$) | Alcoholic<br>controls<br>($n = 29$) | Diabetic<br>controls<br>($n = 40$) |
| --- | --- | --- | --- | --- | --- | --- | --- |
| **Sex (female/male)** | 30/79 | 56/164 | 51/89 | 276/953 | 5/16 | 7/22 | 9/31 |
| **Age (years)** | 64<br>(58-71) | 60<br>(49-69) | 70<br>(52-82) | 45<br>(30-60) | 66<br>(60-74) | 65<br>(61-69) | 62<br>(45-70) |
| **BMI\* (kg/m<sup>2</sup>)** | 21.3<br>(18.0-24.5)<br>Obs. = 99% | 24.2<br>(20.8-27.8)<br>Obs. = 99% | 24.1<br>(20.7-27.7)<br>Obs. = 96% | 23.9<br>(21.4-27.0) | 15<br>(13.1-18.1) | 24.1<br>(22.0-27.5) | 25.8<br>(21.9-31.2) |
| **Acetone\* (‰)** | 0.1<br>(0.1-0.2)<br>Obs. = 73% | 0.3<br>(0.2-0.4)<br>Obs. = 75% | 0.1<br>(0-0.2)<br>Obs. = 14% | 0<br>(0-0.1)<br>Obs. = 1% | 0.1<br>(0-0.1)<br>Obs. = 52% | 0.2<br>(0-0.3)<br>Obs. = 14% | 0.2<br>(0.2-0.3)<br>Obs. = 20% |
| **BHB\* (µg/g)** | 520<br>(385-940)<br>Obs. = 98% | 880<br>(380-1,000)<br>Obs. = 97% | 125<br>(0-278)<br>Obs. = 64% | 0<br>(0-73)<br>Obs. = 6% | 220<br>(165-280)<br>Obs. = 71% | 0<br>(0-298)<br>Obs. = 69% | 63<br>(0-170)<br>Obs. = 72% |
| **Ethanol\* (‰)** | 0<br>(0-0.5) | 0<br>(0-0)<br>Obs. = 99% | 0<br>(0-0.05)<br>Obs. = 25% | 0<br>(0-0.5)<br>Obs. = 99% | 0<br>(0-0)<br>Obs. = 95% | 0.6<br>(0-1.5)<br>Obs. = 90% | 0<br>(0-0)<br>Obs. = 98% |
| **Glucose\* (mmol/L)** | 0.4<br>(0.2-0.7)<br>Obs. = 76% | 33.1<br>(18.3-47.5)<br>Obs. = 85% | 1.0<br>(0.2-3.6)<br>Obs. = 70% | 0.2<br>(0.1-0.5)<br>Obs. = 16% | 0.5<br>(0.3-0.6)<br>Obs. = 71% | 0.8<br>(0.3-3.1)<br>Obs. = 72% | 10.4<br>(2.4-21.9)<br>Obs. = 90% |
| **PMI<sup>1,\*</sup> (days)** | 5.5 (3-6.3)<br>Obs. = 15% | 5 (3.8-8.3)<br>Obs. = 15% | 5.5 (3.3-10.8)<br>Obs. = 24% | 5 (3-7)<br>Obs. = 53% | 6 (6-6)<br>Obs. = 14% | 6 (3-9.5)<br>Obs. = 24% | 5 (3.5-5)<br>Obs. = 28% |
| **PMI<sup>2,\*</sup> (days)** | 8 (6-16.3)<br>Obs. = 84% | 9 (6-13)<br>Obs. = 85% | 8 (6-13)<br>Obs. = 75% | 5 (3-7)<br>Obs. = 47% | 10 (7-13.5)<br>Obs. = 76% | 6.5 (4.3-12)<br>Obs. = 76% | 7 (6-11)<br>Obs. = 72% |

Note: AKA – alcoholic ketoacidosis, DKA – diabetic ketoacidosis. Continuous variables are presented as median and interquartile ranges (Q1-Q3). \* = variable is not observed in all cases. Obs. = percentage of cases with an observed value (if not provided: variable is observed in all cases). BHB > 1,000 is set to upper limit of detection (1,000 µg/g). Glucose < 0.2 is set to 0. Ethanol < 0.1 is set to 0. Postmortem intervals (PMI) are based on the time between the day of sampling and, either day of witnessed death (PMI<sup>1</sup>) or day of last seen alive (PMI<sup>2</sup>).

### Multivariate statistical analyses showed 1,416 significantly different abundant metabolic features between ketoacidosis cases and controls

We first sought to quantify the metabolomic changes related to death by ketoacidosis. To this end, we applied Mann-Whitney U tests with Bonferroni correction, which revealed 1,416 metabolic features with a significant difference in abundance between ketoacidosis cases and controls. Metabolic features were adjusted for PMI, BMI, age, and gender using a metabolite-wise linear modelling and an empirical Bayes moderation of the variances. The obtained p-values were adjusted with a Benjamini-Hochberg false discovery rate correction, and metabolic features with a q-value < 0.05 and an absolute log_2_ fold change (Log_2_ FC) > 1.5 were considered significantly abundant in the volcano plot (Figure 1). After controlling for PMI, BMI, sex, and age, 133 metabolic features were found to be significantly more abundant in ketoacidosis cases compared to controls (red) and 37 were found to be significantly less abundant in ketoacidosis compared to controls. The remaining 4,314 metabolic features were considered not significant (grey). The significantly abundant metabolic features that could be annotated during the feature annotation, are labelled in Figure 1.

To analyse these differences from a systemic perspective, we performed a principal component analysis (PCA). In this analysis, the first and second components captured 7.4% and 4.1% explained variance, respectively (Figure S1). However, distinct patterns were not visible on these principal components, suggesting that supervised machine learning approaches are needed to determine ketoacidosis cases in forensic casework.

### Binary Classification Models Accurately Distinguish Ketoacidosis from Controls

Since there was no clear separation visible on the PCA plot, we employed three different supervised machine learning classification models: Random Forest (RF), Least Absolute Shrinkage and Selection Operator (LASSO), and Support Vector Machine with a linear kernel (SVM) to distinguish the ketoacidosis cases from controls. These three binary classification models were trained on the same training set consisting of 70% of the ketoacidosis cases and 70% of the controls. Additionally, the same test set, consisting of 30% of ketoacidosis cases and 30% of controls, was used to evaluate the three binary classification models. The classification results of the test set, visualised with a confusion matrix, are found in S2A-B for the SVM model and the LASSO model. The classification results of the RF model are found in Figure 2A. Furthermore, the predicted probabilities by the RF model of the ketoacidosis cases and controls in the test set are shown in Figure 2C.I and 2C.II, respectively. The three models had a true positive (TP) rate of 80.9% to 89.4% and a true negative (TN) rate of around 98%. The remaining cases were misclassifications, with a false positive (FP) rate of around 2% and a false negative (FN) rate of 10.6% to 19.1%. An overview of the performance metrics of the three binary classification models can be found in the table in Figure 2B. When looking at the performance metrics, all three models performed similarly (Figure 2B). Thus, all binary classification models performed well in separating ketoacidosis cases and controls.

### Independent Starvation Cases were Predicted as Ketoacidosis

We next investigated whether the models could generalize to correctly predict ketoacidosis beyond their initial training data. To this end, we extracted metabolomics data from known starvation cases (*n* = 21) to use as an independent cohort for model validation. Starvation typically involves low levels of insulin and the elevated metabolism of fat, which may lead to ketoacidosis. Importantly, no starvation cases were used in the training of the models. With the RF model, 17 starvation cases out of 21 (81%) were classified as ketoacidosis (expected 10.5 under the null hypothesis, p_binom_ = 0.003). With the LASSO and SVM models, 18 and 19 of the starvation cases out of 21, respectively, were classified as ketoacidosis (p_binom_ < 0.001 for both models). The predicted probabilities by the binary RF model are shown in Figure 2D. The prediction results of the models are found in S3A-C. Overall, starvation cases were predicted to be more similar to ketoacidosis cases than controls.

### Multinomial Classification Models Separate Ketoacidosis into Subtypes

Knowing that the models could correctly identify ketoacidosis in real-world samples, we investigated if known subtypes of ketoacidosis could be distinguished. This is important as ketoacidosis can have different causes, which cannot always be disentangled with biomarker measurements, like ketone body levels. We used the same three model types (RF, LASSO and SVM) in this multinomial classification between the groups AKA, DKA, hypothermia, and controls. Importantly, the models could predict the correct subclass in the test set with a balanced accuracy of 0.83-0.88 (Figure 2B). The recall in the RF model was 57.6% for AKA and 84.8% for DKA (Fig. 3A-B). The other two models performed similarly and confusion matrices for test set predictions by the LASSO and SVM models can be found in S4.1-2. Figure 3C shows the predicted probabilities of the RF model with the means denoted by black bars.

**Figure 3.**
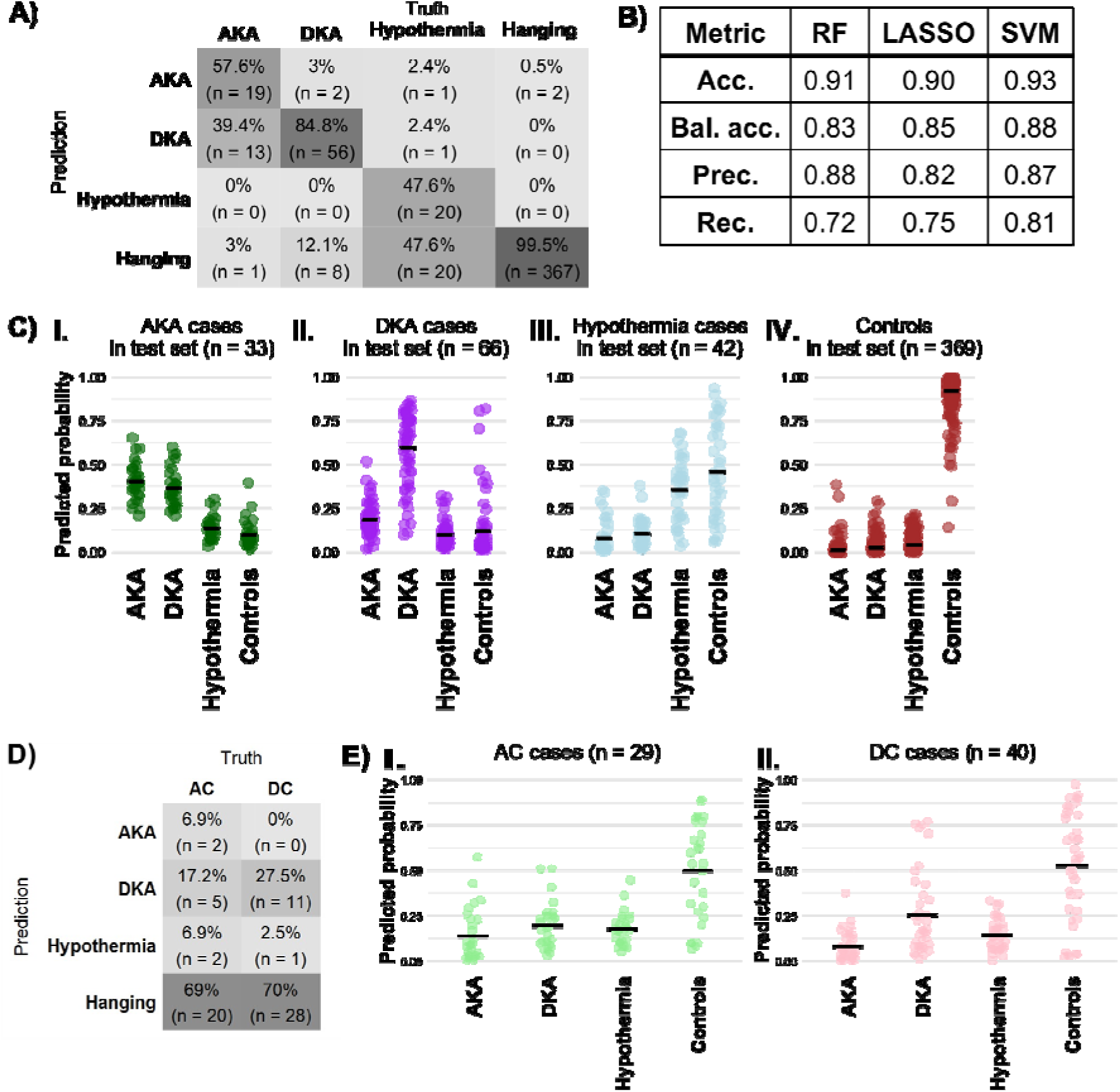
Supervised ML models could distinguish between different ketoacidosis cases and controls. A) Confusion matrix for the multinomial RF model. B) Multi-class performance metrics of the three different multinomial models. C) Predicted probabilities of different classifications of test set data by the multinomial RF model. I: AKA cases, *n* = 33. II: DKA cases, *n* = 66. III: hypothermia cases, *n* = 42. IV: controls (hanging cases), *n* = 369. Means denoted with black bars. D) Predicted classes for the multinomial RF model when tested on AC and DC cases. E) Predicted probabilities of different classifications of AC and DC cases by the multinomial RF model. I: AC cases, *n* = 29. II: DC cases, *n* = 40.

### Proper Independent Cohort Selection is Important

Having found machine learning models to be useful in separating subclasses of ketoacidosis, we investigated to what extent these predictions could generalize. In detail, we sought to test if AKA and DKA were classified based on ketoacidosis-related signals, or metabolomic changes related to alcoholism and diabetes, respectively. To this end we used an independent validation cohort consisting of AC (*n* = 29) and DC (*n* = 40), i.e. individuals with alcohol (AC) or diabetes (DC) indicated as secondary cause of death, that had a main cause of death other than ketoacidosis. The same three models were used for classification of AC and DC cases without retraining. For the RF model (Figure 3D), around 70% of cases in each group (AC and DC) were predicted to be most like controls (hanging cases). However, the LASSO and SVM models struggle to classify AC and DC cases as controls (Figure S5A-B). DC cases were most often classified as controls and after that as DKA cases, indicating that the models partially capture the underlying diabetic pathological condition. These results highlight the importance of study design and highlight that models trained without independent cohorts may overfit to disease-specific metabolic signatures.

### Misclassifications in Binary and Multinomial Classification

To better understand the models, we further examined the misclassified cases in both the binary classification and multinomial classification. The details of FN and FP of each binary classification model are summarized in Table S8, and the FN are visualized using an UpSet plot (Figure S8). When looking closer at the misclassifications by the binary classification models, hypothermia cases are overrepresented. AKA cases, on the other hand, were almost never misclassified by the three models. Furthermore, there were two specific DKA cases and nine hypothermia cases which were misclassified by all three binary classification models.

In multinomial classification, the RF model struggled to predict hypothermia cases, mostly conflating hypothermia cases with controls. Hypothermia cases were included in the ketoacidosis group because the metabolome after hypothermic death closely resembles ketoacidosis, including elevated ketone body levels, due to altered thermogenesis, overall metabolic stress, and energy metabolism changes [5,29,30]. Similarly, elevated BHB levels are present in our hypothermia cases (see Table 1) but are on average lower than the BHB levels in the other ketoacidosis groups (AKA, DKA, and starvation). Still, the BHB levels are considerably higher than in the control groups (hanging, AC, and DC).

The second-most prevalent misclassification was AKA wrongly predicted as DKA. Multinomial classifications of the three models had overlaps for three AKA cases predicted as DKA cases and for 10 hypothermia cases predicted as controls. Furthermore, 11 of the 40 DC cases were classified as DKA cases. Interestingly, among the eight overlapping DC cases that were classified as DKA during multinomial classification, five had diabetic ketoacidosis as a secondary but not primary cause of death. Importantly, nearly all control cases were correctly classified (Fig. 3A-B), indicating high specificity of the model in distinguishing ketoacidosis from controls.

### Clustering of Important Features Reveals Grouping Based on Cause of Death, Not Underlying Conditions

To elucidate the features used in multinomial classification, we extracted the top 25 features that were used in the multinomial RF model training (Figure S6). The clustered median abundances of these features, and the mass over charge and retention times in seconds (in brackets), can be found in Figure 4. Interestingly, control cases were clustered together, and AKA/DKA cases clustered together. Hypothermia cases clustered closer to controls than the other ketoacidosis-related cases. In these top 25 features, there were several named metabolic features, namely: nudifloramide (2PY), N^1^-methyl-4-pyridone-3-carboxamide (4PY), cortisol, glucosamine, and 5,6-indolequinone-2-carboxyilic acid (boxplots can be found in S7).

**Figure 4.**
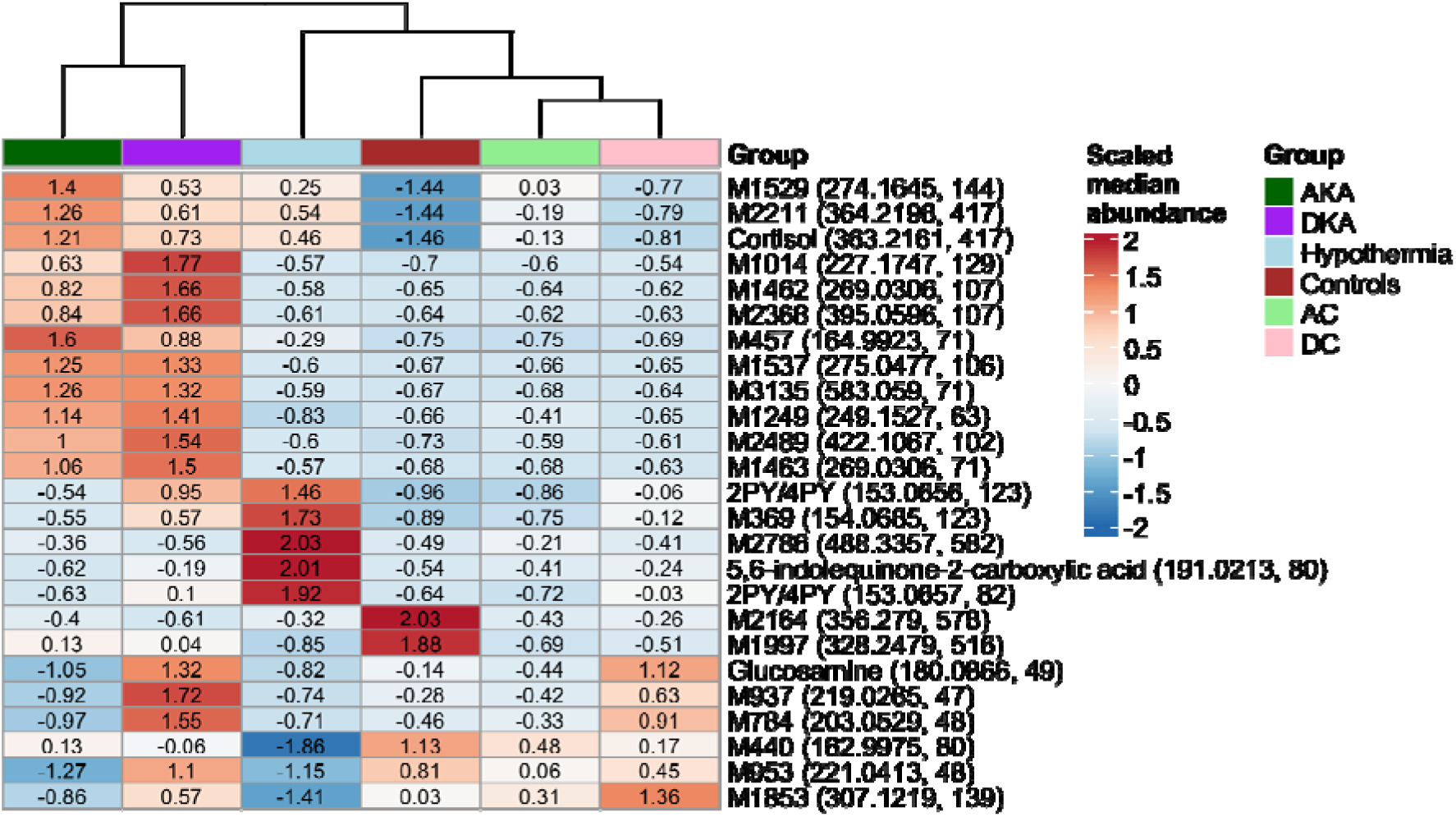
Heatmap of the scaled median abundances of the top 25 important metabolic features. The mass over charge and retention times (in seconds) have been added to each metabolic feature.

## Discussion

This study demonstrates, for the first time in real forensic cases, that supervised machine learning models can accurately detect (balanced accuracy > 0.90) and subtype (balanced accuracy > 0.80) ketoacidosis cases using postmortem metabolomics data. The binary classification model, as trained for the detection of ketoacidosis, could classify unseen starvation cases as more similar to ketoacidosis cases than controls. Taken together, these results show that the integration of supervised ML models and postmortem metabolomics is a valuable tool in forensic decision-making in ketoacidosis-related deaths.

Binary classification models have been used in many different fields to assign cases to one of two categories based on their input data [31]. Most literature related to binary classification models focuses on antemortem characteristics, for example predicting cause [32] or time of death based on clinical data [33]. Nevertheless, Elmsjö et al. (2024) reported a TP rate 92% and TN rate 96% using an Orthogonal Partial Least Squares Discriminant Analysis (OPLS-DA) binary classification model. In the current study, in contrast, we found a higher misclassification rate in hypothermia cases, with most misclassified hypothermia cases instead being classified as controls (hanging cases) (see Figure 3 and S3A-B).

Ward et al. (2024) employed a supervised OPLS-DA multinomial classification model and reported a TP rate of 56% of acidosis cases in the test set, while here we report a higher TP rate of 81% to 89% in binary classification (see Figure 2, S2A-B and S3A-C). Our multinomial models have a lower TP rate in hypothermia cases and a higher TP rate in other ketoacidosis-related cases (AKA and DKA) (see Figure 3 and S5A-B). Furthermore, the classification hanging cases in Ward et al. resulted in a TP rate of 73%, while we report a higher TP rate by all models, both in the binary and multinomial classification of hanging cases (i.e., controls). Even though there is an overlap in forensic cases used in this and the study by Ward et al., we have showed that the models developed here performed better for ketoacidosis detection.

In our study, we could annotate 201 out of the 4,484 features to known metabolites, which limits the mechanistic interpretation of our identified markers. Nevertheless, our computational analyses identified metabolic features associated with ketoacidosis. Among the top 25 metabolic features was glucosamine, which was more abundant in the DKA and DC groups (Figure 4). Hexosamines, like glucosamine, are involved in diabetic physiopathology, like insulin resistance, and can thus be considered a diabetic marker [34–36]. Cortisol has been reported as a biomarker for hypothermia (Elmsjö et al., 2024, Shida et al., 2020). This finding was confirmed in our study where we show that cortisol is higher in all ketoacidosis subtypes compared to controls. Similarly, 2PY/4PY, breakdown products metabolized from excess Vitamin B3 and NAD, was found to be more abundant in hypothermia cases compared to controls, which has also been found by Elmsjö et al. and was found to be upregulated in hypothermic living individuals [38]. Interestingly, Vitamin B3 induced decreases in body temperature in mice, partially due to heat loss due via vasodilation [39]. We also found 5,6-indolequinone-2-carboxyilic acid to be higher in hypothermia. Indolequinones are important in the biosynthesis of melanin [40]. The (potential) role of 5,6-indolequinone-2-carboxyilic acid with regards to cause of death is, however, unknown.

A major strength of this study lies in the dataset itself, which is unique in both size and real-world forensic variability. Unlike controlled clinical datasets, these cases capture authentic forensic complexity, including overlapping pathologies, rare conditions, and diverse metabolic profiles. This heterogeneity mirrors the challenges encountered in routine forensic practice, making the modelling results relevant for practical casework. By training and validating models on such data, we ensure that the predictive performance reflects real-world conditions rather than idealized scenarios, thereby increasing the potential for successful implementation in forensic workflows.

This study has several limitations. In Swedish forensic practice, a forensic pathologist generally assigns a cause of death for every case that undergoes a forensic autopsy. To account for situations where the findings are inconclusive or the underlying mechanism remains uncertain, the cause of death is accompanied by a statement describing the certainty of the diagnosis. This statement has not been available in the present study. Consequently, the ground truth labels used here reflect the forensic pathologist’s best determination, which may not always correspond perfectly to the true cause of death. Additionally, the secondary cause of death may have driven the terminal metabolic state or even be the true cause of death. The dataset reflects a single national forensic system and analytical pipeline, which may limit generalizability to other jurisdictions, instruments, or workflows. We selected three supervised machine learning modelling approaches, while alternative models such as gradient-boosted trees or deep neural networks might better capture nonlinear relationships and improve performance if more data was available. Models were developed based on only metabolomic features without incorporating available covariates (e.g., age, sex, postmortem interval), which could potentially enhance discrimination between groups. The data were collected from authentic autopsy cases, and it is possible that more covariates beyond what is recorded in the autopsy reports affect the metabolomic signal. Nevertheless, the correction for the available covariates, as shown in Fig. 1, showed a considerable signal, with metabolites relevant to ketoacidosis. Hanging cases were selected as controls under the assumption of minimal influence on postmortem metabolomic profiles due to the rapid nature of the process [41]. However, the partial overlap between control cases with hypothermia in important features (Figure 4) indicates that hypothermia and hanging share metabolic patterns. Finally, metabolite identification remained incomplete, with a small fraction of features annotated, limiting mechanistic interpretations of the results.

In conclusion, our study shows that integrating supervised machine learning models with postmortem metabolomics leads to predictive power for detecting and subtyping ketoacidosis-related deaths in real-world forensic cases. Further research is needed to confirm these results and, hopefully, will lead to widespread use of ML models in forensic casework.

## Materials and methods

### Study Population and Data Selection

Autopsy cases admitted from late June 2017 until November 2020 were considered for inclusion in this study. All considered cases are 18 years or older and have undergone toxicological screening using UHPLC-QTOF using femoral blood (*n* = 17,011). Note, Sweden makes use of the International Statistical Classification of Diseases, 9^th^ Revision (ICD-9) [42]. For better specification, the standard codes are sometimes suffixed with a letter according to the Swedish ICD-9 codes, of which some are specific to forensic pathology. Alcoholic ketoacidosis (AKA) and diabetic ketoacidosis (DKA) cases were selected from cases where the main cause of death was determined by the forensic pathologist to be AKA (*n* = 109, ICD-9: 271 or 276) or DKA (*n* = 220, ICD-9: 250), respectively. We also selected hypothermia cases which had hypothermia as main cause of death (*n* = 140, ICD-9: 991). Controls were selected from cases where the main cause of death was typical, mechanical hanging cases, of which the overwhelming majority were suicides (*n* = 1,229, ICD-9: 994). Starvation cases were selected from cases where the main or secondary cause of death was due to nutritional deficiency (*n* = 21, ICD-9: 261, 263 or 269), with or without specified acidosis. Alcoholic and diabetic controls (AC and DC) were selected from cases that had different main causes of death (ICD-9: 348, 410, 411, 422, 428, 431, 432, 441, 800, 801, 806, 807, 808, 820, 852, 864, 958, 965, 977, 980, 986, or 994) and alcohol or diabetes specified as secondary cause of death (AC, *n* = 29; DC, *n* = 40), as determined by the forensic pathologist responsible for the case. This assessment could be based on case circumstances, autopsy findings, or a combination of these. Due to limited sample size, AC and DC were not matched to AKA or DKA cases. In total, there were 7 groups (*N_total_* = 1,788) considered in this study and a demographic overview of the groups can be seen in Table 1.

### Institutional review board statement

This study was approved by the Swedish Ethical Review Authority (dnr: 2019-04530). Since this is a retroactive study, the need for informed consent was waived by the Swedish Ethical Review Authority. All methods were performed in accordance with relevant guidelines and regulations.

### Data acquisition

Each sample was prepared by protein precipitation with the addition of three internal standards: amphetamine-D8, diazepam-D5, and mianserin-D3. All prepared samples were analysed using a UHPLC–ESI–QTOF system. Chromatographic separation was achieved on a C18 column under gradient elution conditions. Mass spectrometric data were acquired in positive ionization mode, with a total run time of 12 minutes per sample. Each sample was required to meet the predefined internal standard acceptance criteria (specific absolute peak areas, retention time deviation within ±0.1 min, and mass accuracy deviation within ±5 ppm). In addition, each analytical batch included a blank whole blood sample fortified with the three internal standards, which was injected at the beginning and at the end of the sequence to monitor system performance. The raw LC/MS data from the selected autopsy cases were exported as mzData files using MassHunter software [21,27].

Further preprocessing and analysis of the data was done in R (version 4.4.1, R Core Team, 2024). First, the postmortem metabolomics data from the cases included in this study were preprocessed using the ‘xcms’ package (version 4.2.3, Benton et al., 2010; Smith et al., 2006; Tautenhahn et al., 2008) and ‘CAMERA’ package (version 1.60.0, Kuhl et al., 2012). In XCMS, the *centWave* algorithm was used for feature detection using the following parameters: Δm/z of 30 ppm, minimum peak width of 3 s, maximum peak width of 30 s, and signal to noise threshold of 3 with the noise variable set to 500. Retention time correction was performed using the *Obiwarp* function and for the grouping an m/z width of 0.05, base width of 3 and minimum fraction of 0.6 were used. More specifically, a feature needed to be present in at least 60% of cases in at least one of the seven groups in this study, for it to be present in the obtained feature peak intensity data. In the end, 4,484 metabolic features could be extracted from the postmortem metabolomics data.

### Data preprocessing and multivariate analysis

The metabolic features data were 0-replaced (feature-wise) and subsequently split into cases for model development and testing (*n* = 1,698) and independent cohorts for model validation (*n* = 90) (Figure 5). The cases for model development and testing consisted of two major groups: ketoacidosis cases (AKA, DKA, hypothermia cases) and controls (hanging cases). 70% of AKA, DKA, hypothermia (**ketoacidosis**, *n* = 469) and hanging (**controls**, *n* = 1,229) were used for model training. The remaining 30% were used for model testing. The independent cohorts used for model validation consisted of starvation cases, AC, and DC, which were never used in model training. Next, the training set, test set, starvation cases, combined AC and DC, were all separately normalized using quantile normalization.

**Figure 5.**
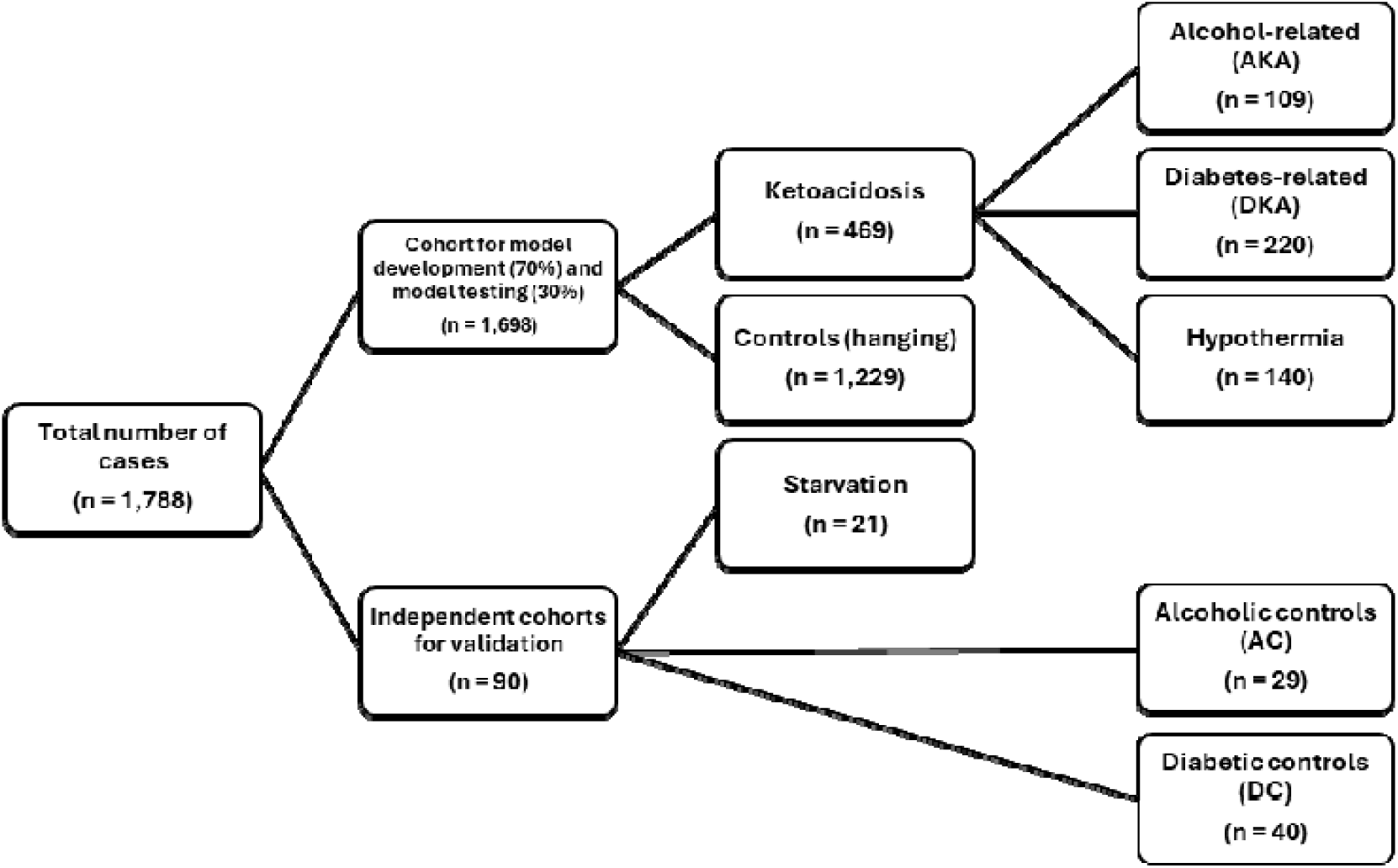
Overview of the data splitting: model development, model testing, and independent cohorts.

### Feature annotation and multivariate analysis

The features were analysed with the “MetaboAnalystR*”* package (version 4.0.0, [48]) to putatively annotate metabolites using the *mummichog* algorithm [49]. Further annotations were manually done based on an inhouse library and online databases (Kyoto Encyclopedia of Genes and Genomes (KEGG) [50], Biochemical, Genetic and Genomic (BiGG) [51], and Human Metabolome Database (HMDB) [52]). In total, 201 out of the 4,484 features could be annotated. Mann-Whitney U tests with a Bonferroni correction were run on each feature to compare abundance levels between ketoacidosis cases and controls.

### Multivariate statistical methods and machine learning models

For data exploration, feature abundances, were visualized with a volcano plot. The feature abundances were adjusted for the covariates PMI, BMI, age and gender, using metabolite-wise linear model fitting with metabolite-wise variance adjustment by the empirical Bayes method, both implemented in the “limma” package [53,54]. Additionally, a principal component analysis (PCA) was run.

For modelling, we employed and compared three supervised machine learning models, namely a random forest (RF) model, a LASSO-penalized logistic regression model (LASSO), and a support vector machines model with a linear kernel (SVM). These three models were used for binary classification and multinomial classification.

The RF model was built with stratified sampling and 500 trees, as implemented in the “randomForest” package [55]. To combat issues due to class imbalance, we used 328 samples from each group in building each tree. For the multinomial RF model, class weights (*classwt* argument) were used instead. Note that the class weights, as implemented in the “randomForest” package, are the priors of the classes, i.e., the number of cases in each group in the training set divided by the total number of cases in the training set. During the LASSO model development, hyperparameter tuning was performed for the value of the tuning parameter λ using a 5-fold cross-validation (λ*_binomial_* = 0.0033 and λ*_multinomial_* = 0.0046). For the multinomial classification, the RF model was run with appropriate class weights based on group size, instead of downsampling the majority group. The binary support vector machines (SVM) model was created using a linear kernel with a cost of constraint violation equal to 10. Radial and polynomial kernels were also tested but did not result in better classification results (not shown). For the multinomial SVM model, again, a linear kernel was chosen with a cost of constraint violation equal to 10.

Lastly, the top 25 important features from the multinomial RF model were extracted based on the mean decrease in accuracy, and the abundances were visualized with a heatmap.

## Data Availability

The data and the corresponding code that support the findings of this study are available in the following repository: https://gitlab.liu.se/ralmo95/project-1-ketoacidosis. Due to legal and ethical considerations, the raw metabolomics data and metadata are not publicly available.

## Author contributions

R.E.C.M., R.M., C.S., H.G., A.E., and E.N. conceived and planned the experiments. R.E.C.M., C.S., and A.E. collected and preprocessed the data. R.E.C.M. performed calculation and modelling analyses with input from all the authors. R.M, C.S., H.G., A.E., and E.N. supervised the work. All authors discussed the results and contributed to the final manuscript.

