## SupplementaryMaterials for "Detecting and Subtyping Ketoacidosis from Metabolomic Patterns in Forensic Casework"

### Supplementary materials

#### S1. PCA plot of ketoacidosis (AKA, DKA, hypothermia) cases and controls (hanging)

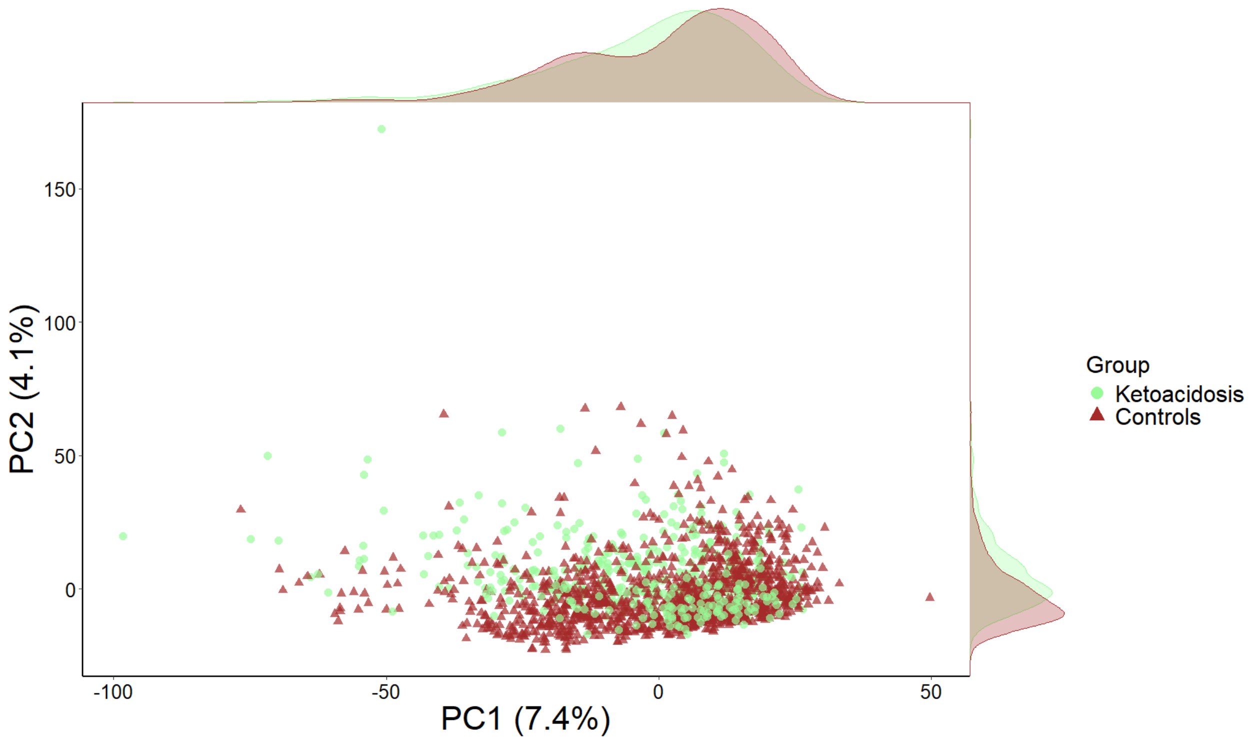


Figure S1. PCA plot of ketoacidosis cases (green) and controls (brown). Principal component 1 has 7.4% explained variance and principal component 2 has 4.1% explained variance.

#### S2. Confusion matrices of binary classification models

**A)**

| Prediction | Truth: **Ketoacidosis** | Truth: **Controls** |
| --- | --- | --- |
| **Ketoacidosis** | 89.4% (n = 126) | 1.6% (n = 6) |
| **Controls** | 10.6% (n = 15) | 98.4% (n = 363) |

**B)**

| Prediction | Truth: **Ketoacidosis** | Truth: **Controls** |
| --- | --- | --- |
| **Ketoacidosis** | 87.9% (n = 124) | 1.9% (n = 7) |
| **Controls** | 12.1% (n = 17) | 98.1% (n = 362) |

Figure S2. Confusion matrices for binary classification models on the test set. A) Confusion matrix for the binary SVM model with a linear kernel B) Confusion matrix for the binary LASSO model.

#### S3. Prediction results: binary classification of starvation cases

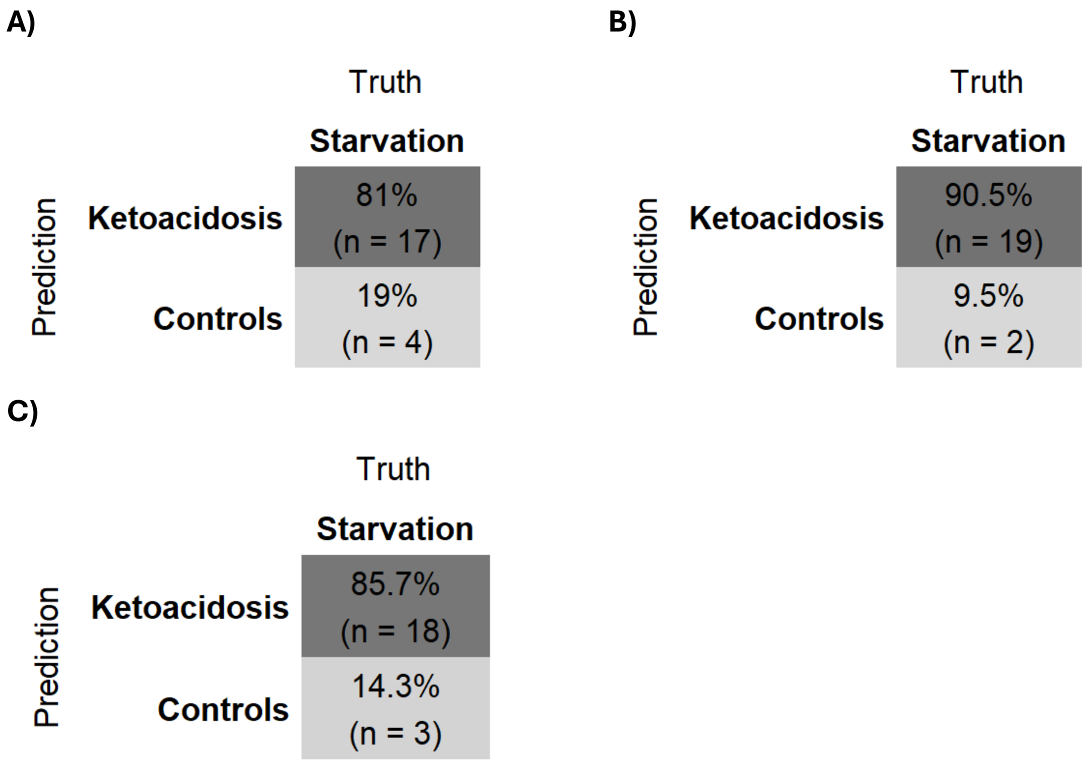


Figure S3. Prediction results of binary classification models tested on starvation cases. B) Prediction results of the binary RF model tested on starvation cases. B) Prediction results of the binary SVM model with a linear kernel tested on starvation cases. C) Prediction results of the binary LASSO model tested on starvation cases.

#### S4. Confusion matrices: multinomial classification models

| Prediction \ Truth | AKA | DKA | Hypothermia | Hanging |
|---|---|---|---|---|
| **AKA** | 63.6% (n = 21) | 4.5% (n = 3) | 2.4% (n = 1) | 0.3% (n = 1) |
| **DKA** | 24.2% (n = 8) | 86.4% (n = 57) | 0% (n = 0) | 0.3% (n = 1) |
| **Hypothermia** | 6.1% (n = 2) | 3% (n = 2) | 73.8% (n = 31) | 0.3% (n = 1) |
| **Hanging** | 6.1% (n = 2) | 6.1% (n = 4) | 23.8% (n = 10) | 99.2% (n = 366) |

Figure S5.1. Confusion matrix for multinomial classification using the SVM model with a linear kernel.

| Prediction \ Truth | AKA | DKA | Hypothermia | Hanging |
|---|---|---|---|---|
| **AKA** | 57.6% (n = 19) | 6.1% (n = 4) | 2.4% (n = 1) | 0% (n = 0) |
| **DKA** | 18.2% (n = 6) | 80.3% (n = 53) | 2.4% (n = 1) | 0.5% (n = 2) |
| **Hypothermia** | 12.1% (n = 4) | 7.6% (n = 5) | 61.9% (n = 26) | 1.1% (n = 4) |
| **Hanging** | 12.1% (n = 4) | 6.1% (n = 4) | 33.3% (n = 14) | 98.4% (n = 363) |

Figure S5.2. Confusion matrix for multinomial classification using the LASSO model.

#### S5. Prediction results: multinomial classification of AC and DC cases

**A)**

| Prediction \ Truth | AC | DC |
|---|---|---|
| **AKA** | 13.8% (n = 4) | 0% (n = 0) |
| **DKA** | 24.1% (n = 7) | 42.5% (n = 17) |
| **Hypothermia** | 10.3% (n = 3) | 7.5% (n = 3) |
| **Hanging** | 51.7% (n = 15) | 50% (n = 20) |

**B)**

| Prediction \ Truth | AC | DC |
|---|---|---|
| **AKA** | 17.2% (n = 5) | 0% (n = 0) |
| **DKA** | 13.8% (n = 4) | 35% (n = 14) |
| **Hypothermia** | 10.3% (n = 3) | 5% (n = 2) |
| **Hanging** | 58.6% (n = 17) | 60% (n = 24) |

Figure S6. Prediction results of binary classification models tested on AC and DC cases. A) Prediction results of the multinomial SVM model tested on AC and DC cases. B) Prediction results of the multinomial LASSO model tested on AC and DC cases.

#### S6. Top 25 important features in multinomial classification (RF model)

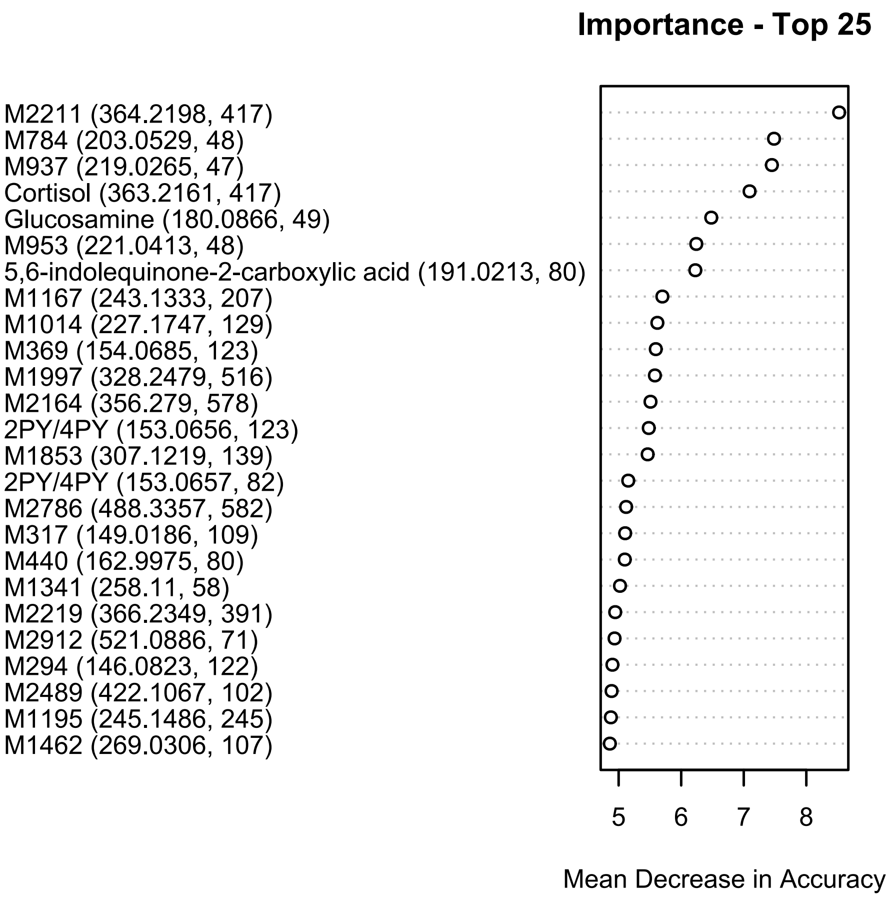


Figure S7. Top 25 important features (in brackets: m/z and retention times in seconds) from the multinomial RF model. Importance based on mean decrease in accuracy.

#### S7. Boxplots of named metabolites in the top 25

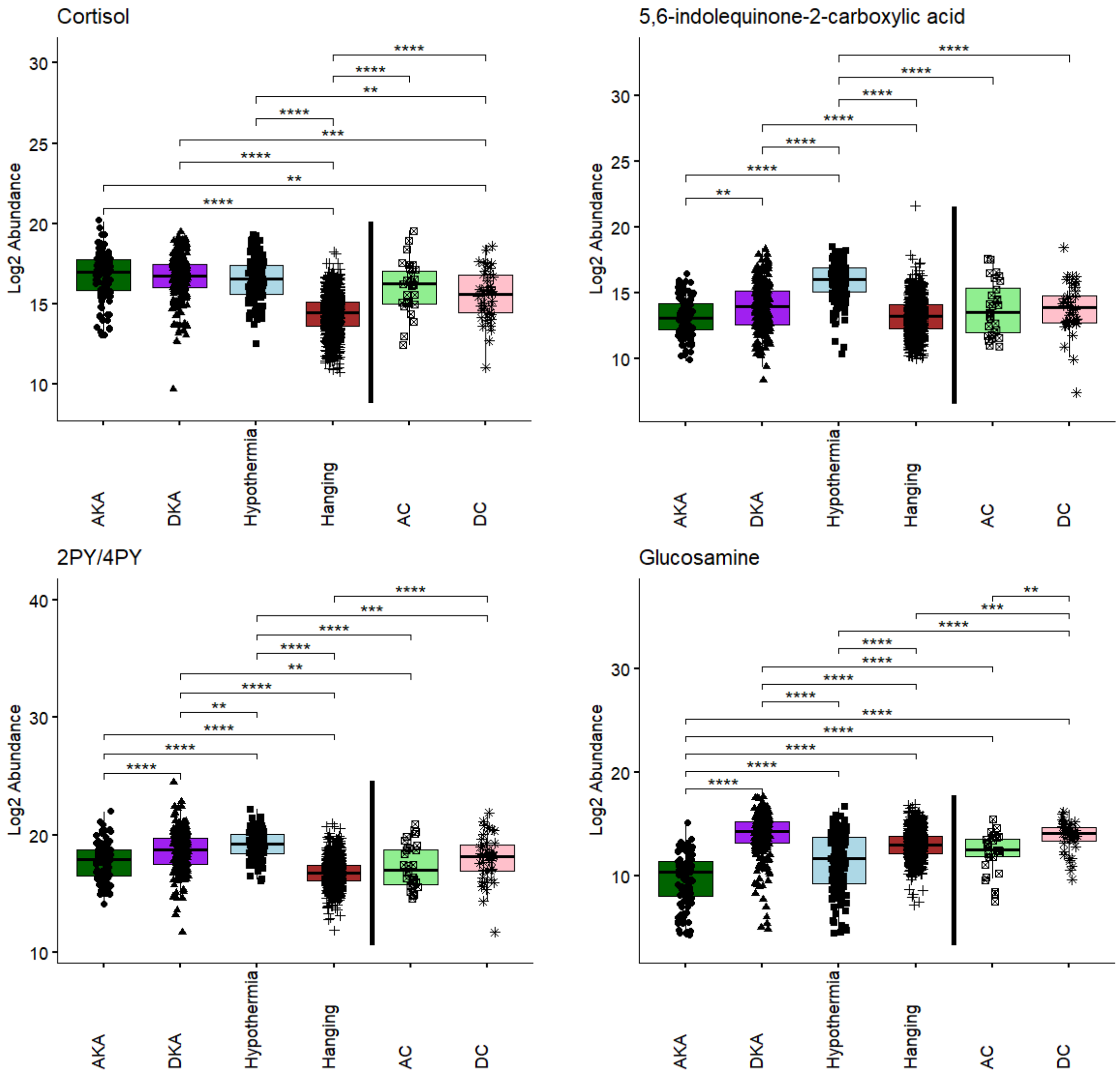


Figure S8. Boxplots showing the log2 abundances of the named metabolites in the top 25. Data were analysed with a Kruskal-Wallis test and pairwise Mann-Whitney U tests, both with Bonferroni correction. Only significant comparisons are shown with significance bars. **p* < 0.05, ***p* < 0.01, ****p* < 0.001, *****p* < 0.0001.

#### S8. Misclassifications in binary classification

Table S8. Summary of FN and FP in binary classification of the test set.

| | RF | LASSO | SVM |
|---|---|---|---|
| **FN: AKA** | 1 | 0 | 1 |
| **FN: DKA** | 4 | 5 | 4 |
| **FN: Hypothermia** | 22 | 12 | 10 |
| **FP** | 4 | 7 | 6 |
| **Total** | **31** | **24** | **21** |

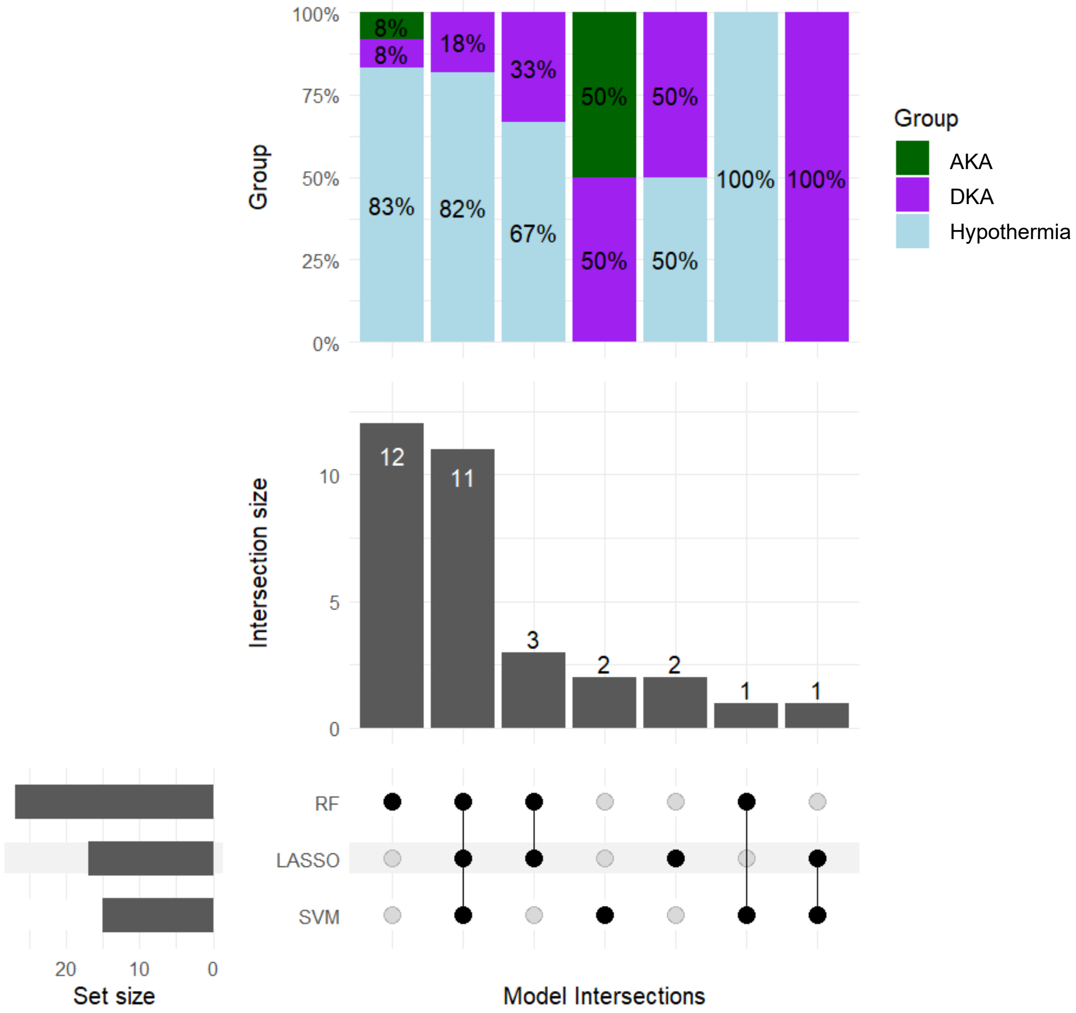


Figure S9. Upset plot showing the overlap and differences in FN cases between the three models.
